# Glioblastoma embryonic-like stem cells exhibit immune-evasive phenotype

**DOI:** 10.1101/2021.12.07.471556

**Authors:** Borja Sesé, Sandra Iñiguez-Muñoz, Miquel Ensenyat-Mendez, Pere Llinàs-Arias, Guillem Ramis, Javier I. J. Orozco, Silvia Fernández de Mattos, Priam Villalonga, Diego M. Marzese

## Abstract

Glioma stem cells (GSCs) are a subset of cells with self-renewal and tumor-initiating capacities that are thought to participate in drug resistance and immune evasion mechanisms in glioblastoma (GBM). Given GBM heterogeneity, we hypothesized that GSCs might also display cellular hierarchies associated with different degrees of stemness. We evaluated a single-cell RNA-seq glioblastoma dataset (*n* = 28) and identified a stem cell population co-expressing high levels of embryonic pluripotency markers, named core glioma stem cells (c-GSCs). This embryonic-like population represents 4.22% ± 0.59 of the tumor cell mass, and pathway analysis revealed an upregulation of stemness and downregulation of immune-associated pathways. Using induced pluripotent stem cell technology, we generated an *in vitro* model of c-GSCs by reprogramming glioblastoma patient-derived cells into induced c-GSCs (ic-GSCs). Immunostaining of ic-GSCs showed high expression of embryonic pluripotency markers and downregulation of antigen presentation HLA proteins, mimicking its tumoral counterpart. Transcriptomic analysis revealed a strong agreement of enriched biological pathways between tumor c-GSCs and *in vitro* ic-GSCs (*κ* = 0.71). Integration of ic-GSC DNA methylation and gene expression with chromatin state analysis of epigenomic maps (*n* = 833) indicated that polycomb repressive marks downregulate HLA genes in stem-like phenotype. Together, we identified c-GSCs as a GBM cell population with embryonic signatures and poor immunogenicity. Genome-scale transcriptomic and epigenomic profiling provide a valuable resource for studying immune evasion mechanisms governing c-GSCs and identifying potential therapeutic targets for GBM immunotherapy.

## Introduction

Glioblastoma (GBM) is the most aggressive primary brain tumor with a median overall survival of 15 months ^1^. Current treatment involves surgical resection followed by concurrent radiotherapy and Temozolomide-based chemotherapy ^2^. However, the systematic development of drug resistance traits is associated with poor prognosis, high recurrence, and high mortality rates. Although immunotherapy (IT) has shown promising results against different solid tumors, phase 3 clinical trials have shown limited efficiency for GBM ^3^. Tumor heterogeneity and phenotypic plasticity combined with a strong immunosuppressive tumor microenvironment remain a challenge for a successful IT in GBM.

GBM contains a subset of glioma stem cells (GSCs), which are thought to possess tumor-initiating capacity and self-renewal potential ^4^. Importantly, GSCs have become a potential target for anti-cancer therapies due to their critical role in tumor aggressiveness and survival^5^, as well as their ability to escape from immune recognition^6^. Nevertheless, purification, expansion, and characterization of GSCs remain a challenging endeavor^7^, potentially in part due to the selective pressure of the culturing conditions favoring specific types of GSC populations. Given the high degree of cellular diversity observed in GBM, we hypothesized that GSCs might also comprise a heterogeneous population with different states of stemness among themselves. Therefore, we sought to find a highly undifferentiated GSC population within the GBM tumor cell mass.

Embryonic stem cells (ESCs) give rise to a wide range of cell types in the adult organism. Interestingly, several groups have reported that ESC-specific gene regulatory signatures are shared among various cancer types^8–11^. Moreover, it has been shown that the ESC core transcription factors OCT4, SOX2, and NANOG are present in GBM tumors, and the upregulation of these factors correlates with poor survival^12,13^. Based on single-cell RNA-seq (scRNA-seq) data from GBM tissues ^14^, we identified a pool of triple-positive GBM cells co-expressing OCT4, SOX2, and NANOG named core GSCs (c-GSCs). Importantly, we observed a significant downregulation of antigen presentation-associated genes in c-GSCs. Using induced pluripotent stem cell (iPSC) technology^15^, we established an *in vitro* cellular model that resembles the tumor c-GSCs. We named these cells induced c-GSCs (ic-GSCs). Additionally, we generated gene expression and DNA methylation (DNAm) maps of ic-GSCs and parental GBM patient-derived cells (GBM-DCs). These data provide hints about the epigenetic and transcriptomic changes involved in immune attenuation mechanisms in ic-GSCs. These findings shed light on the molecular mechanisms underlying immune evasion in GBM.

## Materials and Methods

### Single-cell RNA-seq data analysis

Single-cell RNA-seq (scRNA-seq) data from GBM specimens (n=28) generated by Neftel C et al. was obtained from the Broad Institute Single Cell Portal^14^. All genes with no expression in more than 95% of cells were removed from further analysis. Cells with concomitant high expression of OCT4, SOX2, and NANOG (OSN), defined as the upper quartile expression for each gene, were identified as c-GSCs. Genes with an absolute fold change >0.5 and a Wilcoxon test-based, False Discovery Rate (FDR)-corrected p-value <0.05 were considered differentially expressed genes (DEGs).). Gene Ontology (GO) was used to identify enriched pathways using all DEGs. Radar plots were used to depict HLA-A, -B, and -C (HLA-ABC) levels relative to the OSN profile. R (v.4.0.2) packages *plotly* (v.4.9.4.1), *ggplot2* (v.3.3.2), and *fmsb* (v.0.7.0) were used for data representation.

### Cell lines and culture conditions

GBM-DCs (U3035MG) were obtained from the Human Glioblastoma Cell Culture biobank (Uppsala, Sweden) and cultured under neurobasal media conditions, as previously described^16^. ic-GSCs were cultured on Matrigel-coated dishes (#734-1440, VWR) with mTeSR1 media (#85850, STEMCELL Technologies) as per manufacturer’s instructions.

### Lentiviral production and ic-GSCs derivation

FUW-tetO-hOKMS (Addgene plasmid #51543; RRID:Addgene_51543) was a gift from Tarjei Mikkelsen^17^ and the FUW-M2rtTA (Addgene plasmid #20342; RRID:Addgene_20342) was a gift from Rudolf Jaenisch^18^. GBM-DCs were reprogrammed into ic-GSCs using the iPSC generation protocol as previously described^18^. Briefly, VSVG coated lentiviruses were packaged in 293T cells cultured in mTeSR1 media. FUW-OKMS and FUW-M2rtTA viral supernatants were mixed at a 1:1 ratio, and 0.3 × 10^6^ GBM-DCs were infected four times for 48h. Transduced GBM-DCs were treated with doxycycline 2μg/ml (#72742, STEMCELL Technologies) to induce OKMS expression until ic-GSC colonies appeared after four weeks. Single-cell clone selection was conducted following manual serial dilution in 96-well plates.

### Immunostaining

Cells were fixed in 4% paraformaldehyde for 20 minutes. Immunostaining was performed according to standard protocols. Antibodies were diluted in 1X PBS 0.1% BSA. Primary antibodies and concentration: goat anti-SOX2 1:100 (#LSBio (LifeSpan) Cat#LS-C132162-200, RRID:AB_10833714, R&D), goat anti-OCT4 1:200 (R and D Systems Cat# AF1759, RRID:AB_354975), goat anti-NANOG 1:100 (R and D Systems Cat# AF1997, RRID:AB_355097), and mouse anti-HLA-ABC 1:100 (Thermo Fisher Scientific Cat# MA5-11723, RRID:AB_10985125). Secondary antibodies and concentration: donkey anti-goat IgG (Thermo Fisher Scientific Cat# A32814TR, RRID:AB_2866497) and donkey anti-mouse IgG Alexa Fluor 555 1:500 (Thermo Fisher Scientific Cat# A-31570, RRID:AB_2536180). Nuclei were labeled with DAPI mounting medium (P36962, Thermo Fisher Scientific). Stained cells were analyzed on a Leica TCS SPE confocal microscope (Leica Microsystems, Wetzlar, Germany).

### Cell Sorting

ic-GSCs were dissociated using Accutase (#A1110501, Thermo Fisher Scientific) into single-cell suspension and labeled using double staining with anti-TRA-1-81 APC (#17-8883-42, Thermo Fisher Scientific) and SSEA4 Alexa Fluor 488 (#14-8843-80, Thermo Fisher Scientific). Cells were physically sorted using the FACSAria Fusion (BD Biosciences).

### RNA-sequencing profiling and analysis

Total RNA was extracted from GBM-DCs and TRA-1-81+/SSEA4+ ic-GSCs using the EZNA Total RNA Kit (R6834-01 Omega Bio-Tek). Quality control, mRNA amplification, library preparation, and sequencing were performed at the CRG Genomics Unit (Centre for Genomic Regulation, Spain). Libraries were sequenced 2 * 50+8+16 bp on a HiSeq2500 sequencer (Illumina). RNA-seq data were processed according to Doyle *et al*. protocol^19–21^ (https://training.galaxyproject.org) using the *R/DESeq2* package (v.1.28.1)^22^. RNA-seq counts were transformed to TPM using the R/countToFPKM package (v.1.2.0). The correlation matrix between GBM-DCs and ic-GSCs was plotted using the R/corrplot package (v.0.90). GBM-DC and ic-GSC populations were represented as the mean value of all three GBM-DC replicates and ic-GSC lines, respectively. Genes with an FDR corrected p-value below 10^−10^, and an absolute log_2_ median of ratios over 1.5 were considered DEGs. A representative centroid for the c-GSC population was generated by computing the mean of all genes in c-GSCs. A t-distributed stochastic neighbor embedding (t-SNE) of GBM-DC lines, ic-GSC lines, and a centroid representative of the c-GSC population including all c-GSCs DEGs was plotted using R/Rtsne (v.0.15).

### DNA methylation profiling and analysis

Genomic DNA was extracted from GBM-DCs and TRA-1-81+/SSEA4+ ic-GSCs using the Quick-DNA Microprep Kit (D-3020, Zymo Research). Bisulfite conversion, amplification, and hybridization on the Infinium MethylationEPIC array BeadChip (850K, Illumina) were performed by the Epigenomic Services from Diagenode (Cat nr. G02090000). Raw DNAm data was processed and normalized to beta-values using the R/ChAMP package (v.2.18.3). All probes containing repetitive elements or single nucleotide polymorphisms (SNPs) were removed from downstream analyses. DNAm data was processed and normalized to beta-values using the R/*ChAMP* package (v.2.18.3). All probes containing repetitive elements or single nucleotide polymorphisms (SNPs) were removed from downstream analyses. DNAm levels were standardized by dividing the value of each probe by the mean beta-value in each sample. All probes with differences in beta-value >1 and p-value ≤0.1 were considered differentially methylated sites (DMS).

Promoter location (EPDnew Promoters)^23^ was downloaded from UCSC Genome Browser Table^24,25^. All probes located at ±1.5kb of a transcription start site were used to quantify the DNAm levels in gene promoters. The correlation matrix between GBM-DCs and ic-GSCs was plotted using the R/corrplot package (v.0.90). OSN ChIP-seq peaks were acquired from the NCBI-GEO repository (GSE61475)^26^. OSN binding sites (OSNbs) were obtained using bedtools Intersect Intervals (Galaxy v.2.30.0.0). DNAm levels at ±10kb from OSNbs were represented using deepTools (Galaxy v.2.30.0.0). Representative GBM-DC and ic-GSC populations were represented as the mean value of all three GBM-DC replicates and ic-GSC lines, respectively. The chromatin states have been analyzed using the Epilogos visualization model (https://epilogos.altius.org/) based on epigenomic data sets across 833 biospecimens^27^.

## Data availability

All data generated for this manuscript are available upon request to the corresponding authors.

## Results and Discussion

### Identification and characterization of core-glioma stem cells

The idea that tumors originate from embryonic-like cells is a longstanding hypothesis originally conceived in the 19^th^ century after observing histologic similarities between cancer and embryonic tissue under the microscope ^28^. Over the years, mounting evidence has led researchers to theorize that tumors may arise from tissue-specific stem cells that undergo malignant transformations ^29^. In GBM, the isolation and characterization of GSCs have been carried out by culturing primary tumor cells under neural stem cells serum-free media conditions^30^. However, there have been inconsistencies concerning the GSC model due to the lack of consensus on culture methods and specific GSC markers used among different research groups ^31^. Based on the original concept of the embryonal origin of cancer, we analyzed publicly-available scRNA-seq data from GBM patients (n=28) ^14^ searching for undifferentiated cells harboring ESC-like signatures within the tumor mass. By pooling together all GBM patient cells (7,930 cells), our analyses revealed the presence of 339 cells (4.27%) with the concurrent expression of the core ESC pluripotency markers OSN (Fig. 1A, Link: https://coregliomastemcells.github.io/Fig1a.html). This population, described hereinafter as core GSC (c-GSC), was found in 26 out of 28 patients (92.8%), with an average presence of 4.22% ± 0.59 (Fig. 1B).

**Fig. 1.**
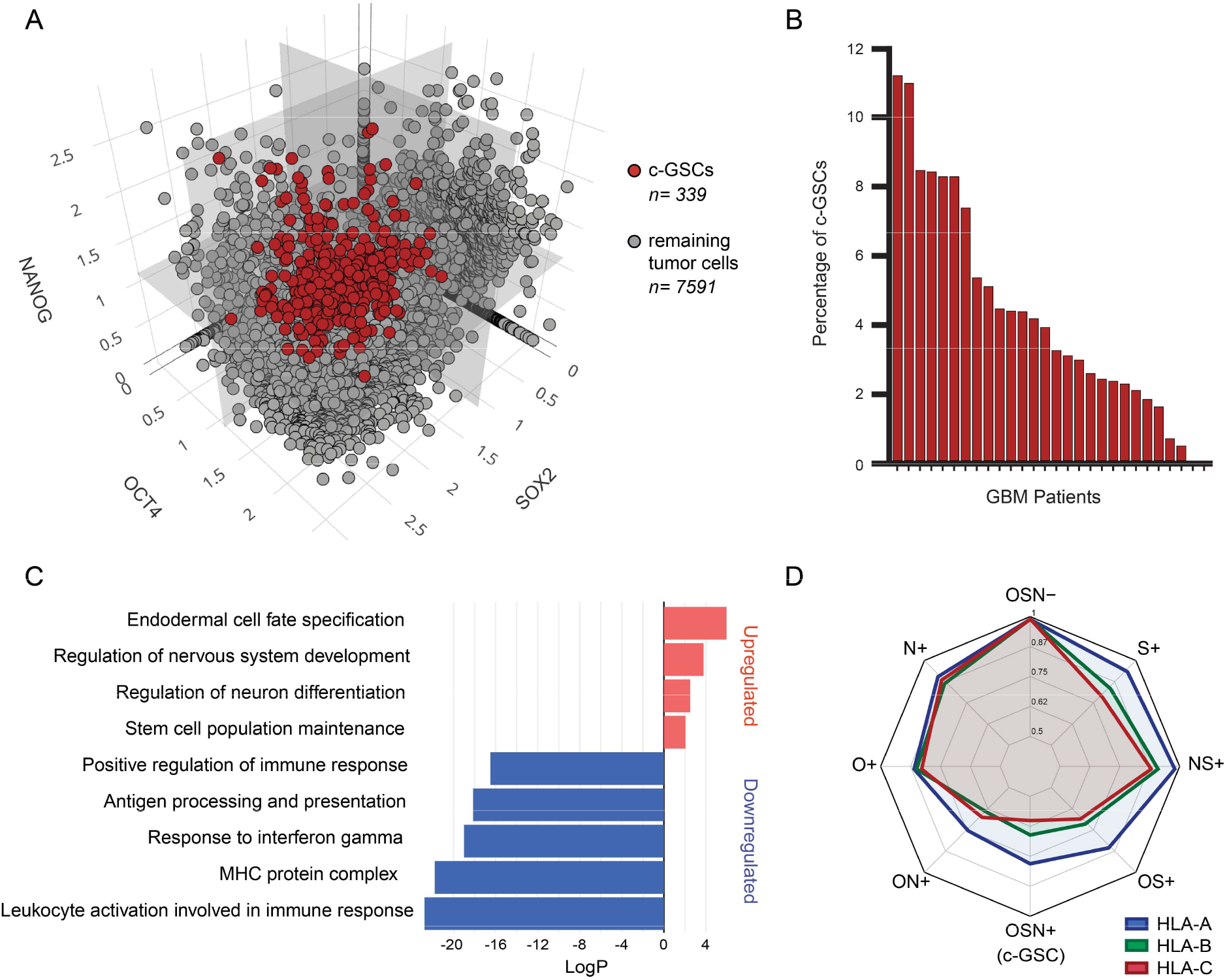
Core glioma stem cell population in glioblastoma samples. (A) Three-dimensional scatter plot of GBM single-cells (n = 7,930) from 28 patients based on the expression levels of OCT4 (x-axis), NANOG (y-axis), and SOX2 (z-axis). Red dots represent c-GSCs co-expressing OCT4, SOX2, and NANOG (n = 339). Gray dots represent the remaining cells from tumor bulk (n = 7,591). An interactive version of this data can be found: https://coregliomastemcells.github.io/Fig1a.html. (B) Bar plot showing the percentage of c-GSCs in each GBM patient. (C) GO analysis of biological pathways upregulated (red) and downregulated (blue) in c-GSCs. (D) Radar plot representing the expression levels of HLA-A, -B, and -C genes on different GBM single-cell populations based on the expression of OCT4, SOX2, and NANOG (OSN).

One way tumors escape from immune recognition is through the downregulation of HLA molecules. Several reports have identified a defect in HLA class I in gliomas associated with immune evasion mechanisms^32,33^. Gene Ontology (GO) analysis revealed a significant downregulation of immune response in c-GSCs, mainly associated with antigen presentation pathways, together with an upregulation of stemness and lineage specification pathways (Fig. 1C). In concordance with this observation, by measuring HLA-A, -B, and -C (HLA-ABC) expression in different tumor bulk cell populations ranging from triple-negative (OSN-) to triple-positive (OSN+), we observed a gradual downregulation of HLA-ABC genes towards the emergence of the c-GSC (OSN+) phenotype (Fig. 1D). These findings led us to speculate that this c-GSC population could have a critical role in GBM tumor growth and invasion, tumor immune-evasion, and ultimately relapse after treatment. It would be important to investigate if the unsuccessful IT is associated with a higher percentage of c-GSCs in the tumor mass. Despite the negative results obtained from clinical trials, multiple efforts are being made to develop novel strategies for GBM IT, including neoantigen vaccines, oncolytic viruses, CAR-T cells, and immune-checkpoint inhibitors^34,35^. Thus, targeting c-GSCs might help elucidate GBM immune evasion-related molecular mechanisms and may provide potential alternatives to improve IT outcomes.

### Reprogramming of glioblastoma-derived cells mimics the core-glioma stem cell phenotype in vitro

A challenge of the current study was the difficulty to isolate and expand c-GSCs *in vitro*, mainly due to sample accessibility and lack of standardized protocols. While these cells represent 4.22% ± 0.59 and are present in 92.8% of the GBM patients, prior GSC studies may have missed the existence of these embryonic-like cells due to the culturing conditions. We used iPSC technology to reconstitute a c-GSC model in cell culture to overcome this. The generation of iPSCs aims to reprogram somatic cells into a pluripotent state, similar to ESCs from the early embryo^15^. Our goal was to reprogram GBM cells and generate cells with a c-GSC-like phenotype that we named ic-GSCs.

GBM-DCs were reprogrammed into induced c-GSCs (ic-GSCs) by ectopic expression of pluripotency-related transcription factors OCT4, KLF4, c-MYC, and SOX2 (Fig. 2A). Three independent ic-GSC lines (ic-GSC#1, ic-GSC#3, and ic-GSC#7) were generated from single-cell clones following serial dilutions and expanded into stable cell lines in the absence of doxycycline. To identify c-GSC molecular signatures in ic-GSCs, we analyzed the expression levels of OSN and HLA-ABC by immunostaining (Fig. 2B). Parental GBM-DCs were positive for SOX2 and HLA-ABC, whereas OCT4 and NANOG remained absent. On the other hand, in concordance with c-GSCs, all ic-GSC lines showed the expression of the three pluripotency-associated markers OSN, along with a strong downregulation of HLA levels. Pluripotent ESCs and iPSCs are immune-privileged by presenting the low expression of HLA class I genes^36,37^, which is associated with an immunological tolerance mechanism of the early embryo to avoid immune rejection by the host^38^. A similar mechanism could occur during tumorigenesis, where the tumor may take advantage of immune-privileged undifferentiated cells to escape from immune system recognition.

**Fig. 2.**
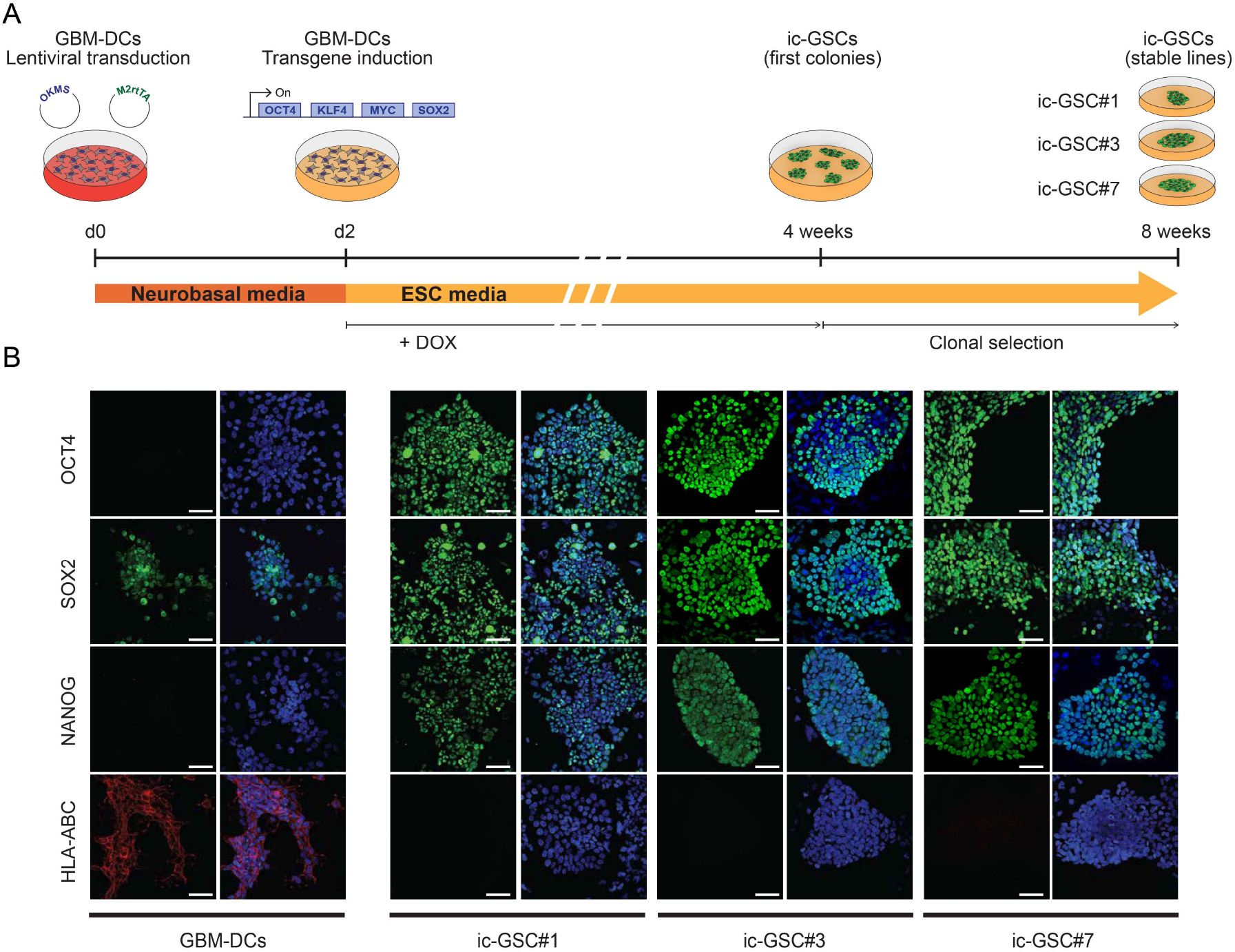
Characterization of *in vitro* induced core glioma stem cell model generated from glioblastoma patient-derived cells. (A) Timeline depicting the reprogramming of GBM-DCs into ic-GSC stable lines. (B) Immunostaining of OCT4, SOX2, NANOG, and HLA-ABC in GBM-DCs and three independent ic-GSC clones: ic-GSC#1, ic-GSC#3, and ic-GSC#7. Scale barrs correspond to 50 μm.

### Gene expression analysis of induced core-glioma stem cells

To validate the reproducibility of ic-GSC lines (ic-GSC#1, ic-GSC#3, ic-GSC#7) and GBM-DC replicates (GBM-DC1, GBM-DC2, GBM-DC3), we measured the correlation among gene expression profiles. We observed that each sample type clustered in two distinct groups (Fig. 3A). In addition to our observations by immunostaining, HLA-ABC showed significant downregulation in ic-GSCs (FDR corrected p <0.0001) compared to the parental GBM-DCs. Then, we performed a t-SNE representation of the three ic-GSC lines, the three GBM-DC replicates, and a centroid representative of the c-GSC population generated from the GBM tumor bulk scRNA-seq data. Our analysis showed clear segregation of all ic-GSC lines from the parental GBM-DC lines and an approximation to the c-GSC population (Fig. 3B). This suggests that reprogramming GBM-DCs into ic-GSCs resulted in a distinct cellular state that resembles the c-GSC gene expression program. To further validate our ic-GSC model, we compared transcriptomic-level profiles of ic-GSC and c-GSC populations (Fig. 3C). We observed that ic-GSCs showed a strong agreement between differentially active gene pathways with c-GSCs, including significant upregulation of stemness and downregulation of immune-associated pathways (OR: 358.20, *κ*=0.71, p <0.0001). Our findings indicate that the transcriptome reprogramming underwent by ic-GSCs *in vitro* recapitulates the gene expression program of c-GSCs from tumor bulk.

**Fig. 3.**
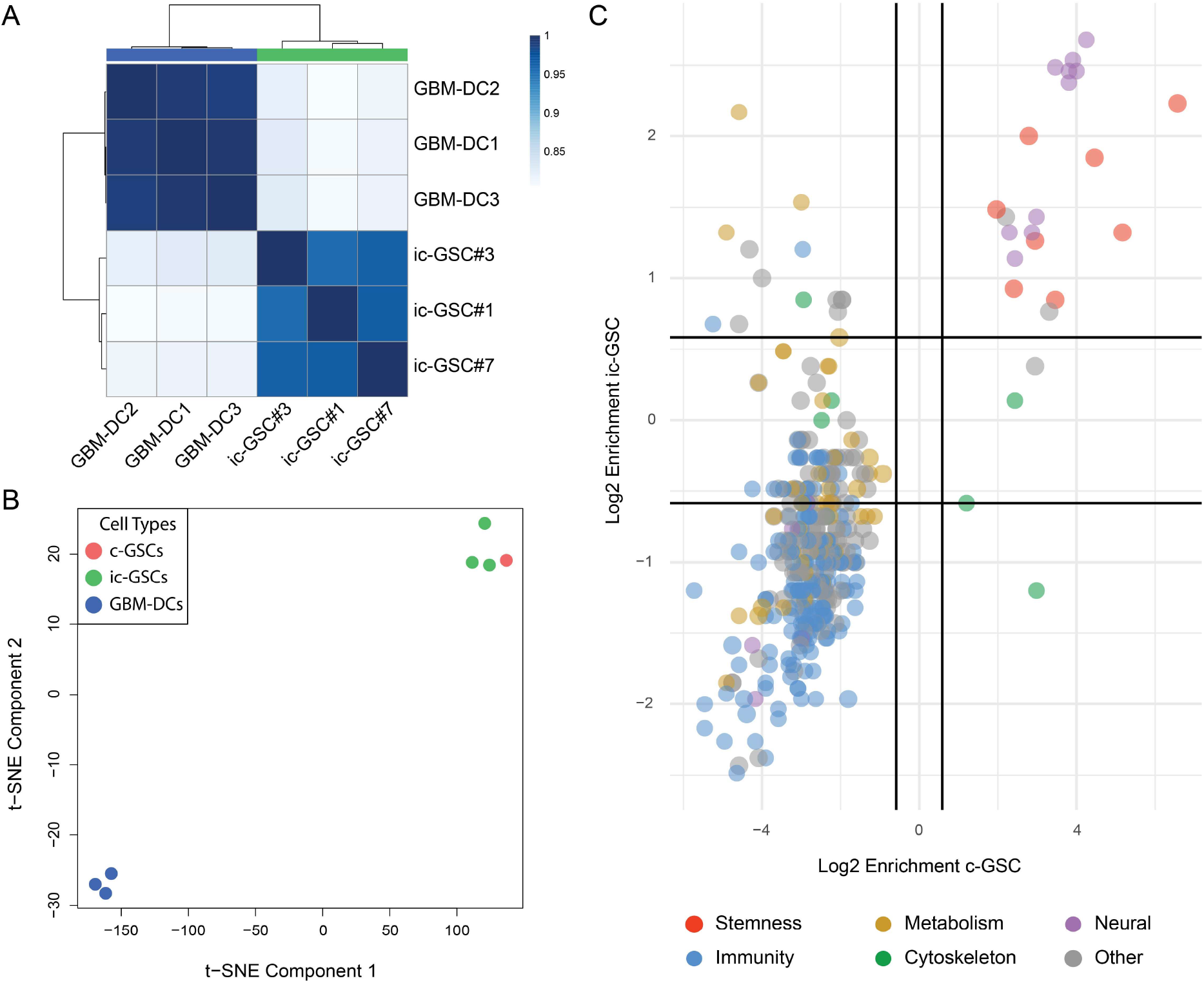
Transcriptomic analysis of induced core glioma stem cells. (A) Correlation matrix for three GBM-DC replicates and three ic-GSC lines based on gene expression profiles. (B) t-SNE plot of three GBM-DC replicates (blue), three ic-GSC lines (green), and a representative centroid of the c-GSC population (red) using gene expression profiles (C) Scatter plot of enriched pathways in representative ic-GSC and c-GSC populations and colored based on their biological network classification. GBM-DC replicates: GBM-DC1, GBM-DC2, and GBM-DC3; ic-GSC lines: ic-GSC#1, ic-GSC#3, and ic-GSC#7.

### Epigenetic changes behind the ic-GSC transcriptome reprogramming

In addition to changes in gene expression, we investigated epigenetic marks potentially involved with the c-GSC aggressive phenotype. We generated DNAm maps from GBM-DCs and ic-GSCs to identify epigenetic changes involved in ic-GSC reprogramming. Similar to the transcriptomic analysis, the genome-wide DNAm showed a significant shift between ic-GSC lines and GBM-DC replicates (Fig. 4A). As expected, we observed that gene regulatory elements binding OSN showed significantly reduced DNAm levels, indicating an active involvement of these elements in ic-GSCs reprogramming (Fig. 4B). Integration of RNA-seq and DNAm data revealed that downregulation of the HLA-C gene correlated with increased DNAm levels at its promoter region (Fig. 4C). In contrast, DNAm at HLA-A and HLA-B promoter regions remained unchanged.

**Fig. 4.**
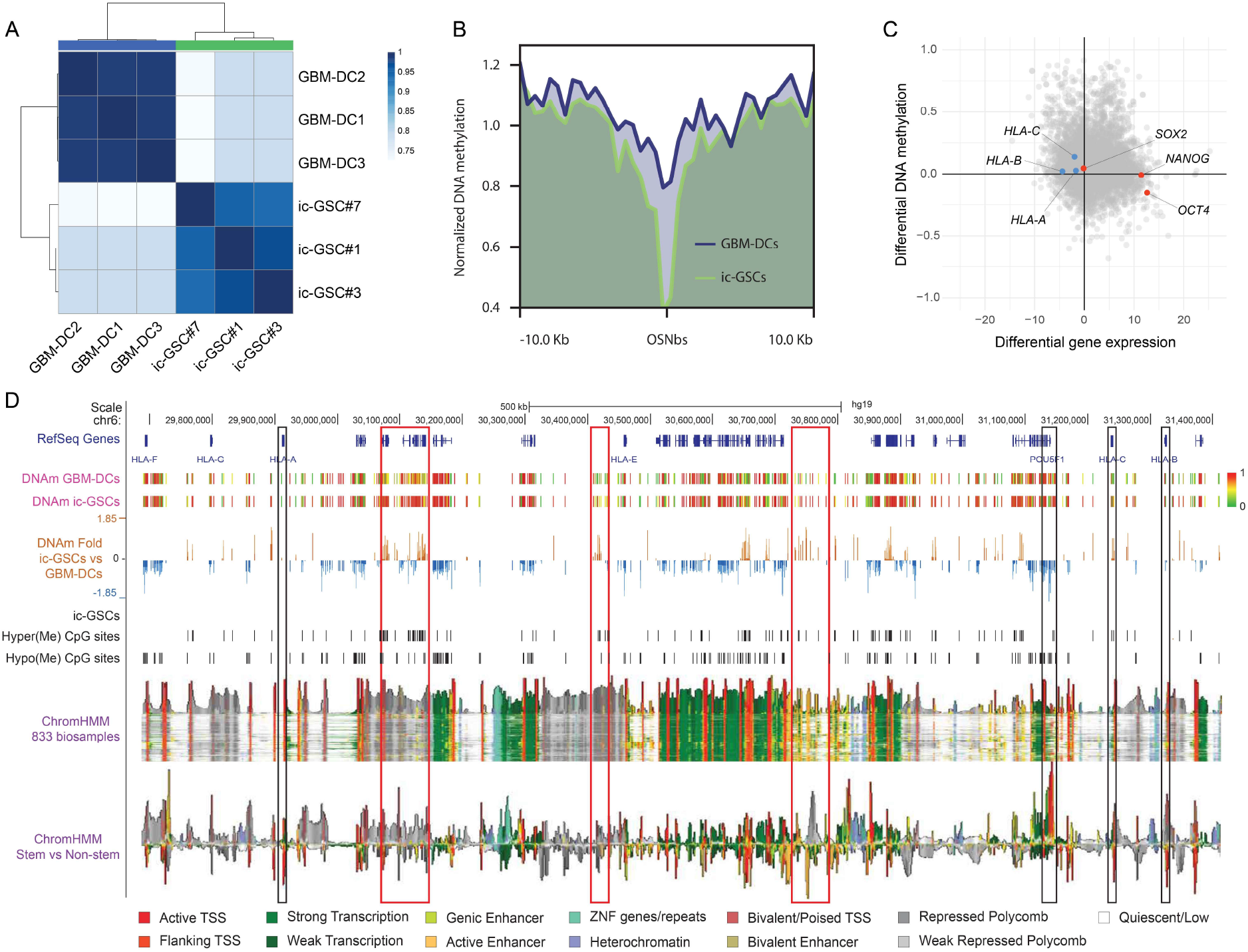
Epigenomic analysis of induced core glioma stem cells. (A) Correlation matrix for three GBM-DC replicates and three ic-GSC lines based on DNAm profile. (B) Profile plot representing the DNAm levels of GBM-DCs (blue) and ic-GSCs (green) around (±10kb) consensus OCT4, SOX2, and NANOG binding sites (OSNbs). (C) Starburst plot including DMS (y-axis) and DEG (x-axis) in representative ic-GSC and c-GSC populations, highlighting stem-related factors OCT4, SOX2, and NANOG (red) and immune-associated antigen presentation genes HLA-A, -B, and -C (blue) (D) Representation of a 1.7Mb genomic region (chr6:29,685,442-31,426,907) encoding HLA-ABC genes and genes relevant for ic-GSC reprogramming. The view includes RefSeq Genes from NCBI, DNAm levels in GBM-DCs and ic-GSCs, DNAm fold change in ic-GSCs vs. GBM-DCs, hyper- and hypomethylated CpG sites in ic-GSCs, chromatin state (ChromHMM) in 833 biosamples, and chromatin state in stem versus non-stem biospecimens. HLA-ABC and OCT4 (*POU5F1*) promoter regions are represented as black rectangles. Hypermethylated regions in ic-GSCs are represented as red rectangles.

On the other hand, endogenous SOX2 and NANOG genes presented no differential DNAm at their promoters. In contrast, the endogenous OCT4 promoter region showed reduced DNAm levels. These findings indicate that DNAm alone does not appear to be the main mechanism for HLA-ABC downregulation, and other epigenetic mechanisms might be involved. To identify potential cis and trans gene regulatory elements whose activation may impact HLA-ABC regulation in c-GSCs, we evaluated the DNAm levels on a 1.7 Mb genomic region (chr6:29,685,442-31,426,907) encoding a cluster of HLA class I genes (from HLA-A to HLA-F; Fig. 4D). Our analysis confirmed the slight DNAm variation at the HLA-ABC promoter regions and a significant hypomethylation of the OCT4 (*POU5F1*) gene (Fig. 4D black rectangles). However, we identified regional changes in DNAm beyond the gene promoter regions. Therefore, we analyzed the chromatin states of the entire region based on a public compendium of epigenomic maps across 833 specimens ^27^. We observed that regions with significant hypermethylation in ic-GSCs overlapped chromatin segments with variable states (Fig. 4D red rectangles). Depending on the cell type and state, these regions can act as enhancer elements (light green and orange pixels), be repressed (grey pixels), or remain quiescent (white pixels).

We compared chromatin states between stem and non-stem samples to evaluate associations with the ic-GSC phenotype. This analysis showed a decrease in active transcription start sites (red pixels) marks in HLA-ABC genes, along with an increment of active TSS in the OCT4 (*POU5F1*) gene. More importantly, hypermethylated regions in ic-GSCs showed a clear increment in polycomb repressed and bivalent poised marks, a phenomenon that was extended to the *HLA-B* and *HLA-C* promoter regions in stem-related samples. A similar phenomenon has recently been observed by Chaligne et al.^39,^ where polycomb activity may establish bivalent domains at hypomethylated promoters in GBM stem-like cells. Accordingly, Burr et al. ^40^ reported that polycomb maintains transcriptional repression of HLA class I genes, keeping them poised by bivalent histone modifications in ESC and cancer cell lines. Thus, the embryonic traits of c-GSCs may lead an epigenetic reprogramming that results in the silencing of HLA genes under a poised state to achieve an immune-evasive phenotype.

Our findings support the existence of a GSC population with embryonic-like features and attenuated immune phenotype. The ic-GSC model provides a valuable in vitro system to study the molecular mechanisms governing c-GSCs immune evasion and may set the basis to improve the current GBM patient’s response to IT.

## References

1. Stupp, R. et al. Radiotherapy plus Concomitant and Adjuvant Temozolomide for Glioblastoma. N Engl J Med 352, 987–996 (2005).

2. Gallego, O. Nonsurgical treatment of recurrent glioblastoma. Curr. Oncol. 22, 273 (2015).

3. Narita, Y. et al. A randomized, double-blind, phase III trial of personalized peptide vaccination for recurrent glioblastoma. Neuro-Oncology 21, 348–359 (2019).

4. Gimple, R. C., Bhargava, S., Dixit, D. & Rich, J. N. Glioblastoma stem cells: lessons from the tumor hierarchy in a lethal cancer. Genes Dev. 33, 591–609 (2019).

5. Zheng, Z.-Q. et al. Nestin+/CD31+ cells in the hypoxic perivascular niche regulate glioblastoma chemoresistance by upregulating JAG1 and DLL4. Neuro-Oncology 23, 905–919 (2021).

6. Alvarado, A. G. et al. Glioblastoma Cancer Stem Cells Evade Innate Immune Suppression of Self-Renewal through Reduced TLR4 Expression. Cell Stem Cell 20, 450-461.e4 (2017).

7. Pollard, S. M. et al. Glioma Stem Cell Lines Expanded in Adherent Culture Have Tumor-Specific Phenotypes and Are Suitable for Chemical and Genetic Screens. Cell Stem Cell 4, 568–580 (2009).

8. Reya, T., Morrison, S. J., Clarke, M. F. & Weissman, I. L. Stem cells, cancer, and cancer stem cells. Nature 414, 105–111 (2001).

9. Ben-Porath, I. et al. An embryonic stem cell–like gene expression signature in poorly differentiated aggressive human tumors. Nat Genet 40, 499–507 (2008).

10. Wong, D. J. et al. Module Map of Stem Cell Genes Guides Creation of Epithelial Cancer Stem Cells. Cell Stem Cell 2, 333–344 (2008).

11. Kim, J. et al. A Myc Network Accounts for Similarities between Embryonic Stem and Cancer Cell Transcription Programs. Cell 143, 313–324 (2010).

12. Bradshaw, A. et al. Cancer Stem Cells in Glioblastoma Multiforme. Front. Surg. 3, (2016).

13. Elsir, T. et al. A study of embryonic stem cell-related proteins in human astrocytomas: Identification of Nanog as a predictor of survival. Int. J. Cancer 134, 1123–1131 (2014).

14. Neftel, C. et al. An Integrative Model of Cellular States, Plasticity, and Genetics for Glioblastoma. Cell 178, 835-849.e21 (2019).

15. Takahashi, K. & Yamanaka, S. Induction of Pluripotent Stem Cells from Mouse Embryonic and Adult Fibroblast Cultures by Defined Factors. Cell 126, 663–676 (2006).

16. Xie, Y. et al. The Human Glioblastoma Cell Culture Resource: Validated Cell Models Representing All Molecular Subtypes. EBioMedicine 2, 1351–1363 (2015).

17. Cacchiarelli, D. et al. Integrative Analyses of Human Reprogramming Reveal Dynamic Nature of Induced Pluripotency. Cell 162, 412–424 (2015).

18. Hockemeyer, D. et al. A Drug-Inducible System for Direct Reprogramming of Human Somatic Cells to Pluripotency. Cell Stem Cell 3, 346–353 (2008).

19. Afgan, E. et al. The Galaxy platform for accessible, reproducible and collaborative biomedical analyses: 2018 update. Nucleic Acids Research 46, W537–W544 (2018).

20. Batut, B. et al. Community-Driven Data Analysis Training for Biology. Cell Systems 6, 752-758.e1 (2018).

21. Doyle, M., Phipson, B. & Dashnow, H. 1: RNA-Seq reads to counts (Galaxy Training Materials). https://training.galaxyproject.org/training-material/topics/transcriptomics/tutorials/rna-seq-reads-to-counts/tutorial.html Online; accessed Thu Sep 30 2021. (2021).

22. Love, M. I., Huber, W. & Anders, S. Moderated estimation of fold change and dispersion for RNA-seq data with DESeq2. Genome Biol 15, 550 (2014).

23. Dreos, R., Ambrosini, G., Groux, R., Cavin Périer, R. & Bucher, P. The eukaryotic promoter database in its 30th year: focus on non-vertebrate organisms. Nucleic Acids Res 45, D51–D55 (2017).

24. Karolchik, D. The UCSC Table Browser data retrieval tool. Nucleic Acids Research 32, 493D –496 (2004).

25. Kent, W. J. et al. The Human Genome Browser at UCSC. Genome Research 12, 996–1006 (2002).

26. Tsankov, A. M. et al. Transcription factor binding dynamics during human ES cell differentiation. Nature 518, 344–349 (2015).

27. Boix, C. A., James, B. T., Park, Y. P., Meuleman, W. & Kellis, M. Regulatory genomic circuitry of human disease loci by integrative epigenomics. Nature 590, 300–307 (2021).

28. Sell, S. Stem cell origin of cancer and differentiation therapy. Critical Reviews in Oncology/Hematology 51, 1–28 (2004).

29. Wicha, M. S., Liu, S. & Dontu, G. Cancer Stem Cells: An Old Idea—A Paradigm Shift. Cancer Res 66, 1883–1890 (2006).

30. Lee, J. et al. Tumor stem cells derived from glioblastomas cultured in bFGF and EGF more closely mirror the phenotype and genotype of primary tumors than do serum-cultured cell lines. Cancer Cell 9, 391–403 (2006).

31. Zhang, L. et al. The necessity for standardization of glioma stem cell culture: a systematic review. Stem Cell Res Ther 11, 84 (2020).

32. Wischhusen, J., Friese, M. A., Mittelbronn, M., Meyermann, R. & Weller, M. HLA-E Protects Glioma Cells from NKG2D-Mediated Immune Responses In Vitro: Implications for Immune Escape In Vivo. J Neuropathol Exp Neurol 64, 523–528 (2005).

33. Feng, E. et al. Correlation of alteration of HLA-F expression and clinical characterization in 593 brain glioma samples. J Neuroinflammation 16, 33 (2019).

34. Majc, B., Novak, M., Kopitar-Jerala, N., Jewett, A. & Breznik, B. Immunotherapy of Glioblastoma: Current Strategies and Challenges in Tumor Model Development. Cells 10, 265 (2021).

35. Guo, X., Wang, S., Wang, Y. & Ma, W. Anti-PD-1 plus anti-VEGF therapy in multiple intracranial metastases of a hypermutated, IDH wild-type glioblastoma. Neuro-Oncology 23, 699–701 (2021).

36. Li, L. Human Embryonic Stem Cells Possess Immune-Privileged Properties. Stem Cells 22, 448–456 (2004).

37. Pick, M., Ronen, D., Yanuka, O. & Benvenisty, N. Reprogramming of the MHC-I and Its Regulation by NF*κ*B in Human-Induced Pluripotent Stem Cells. STEM CELLS 30, 2700–2708 (2012).

38. Gaunt, G. & Ramin, K. Immunological Tolerance of the Human Fetus. Am J Perinatol 18, 299–312 (2001).

39. Chaligne, R. et al. Epigenetic encoding, heritability and plasticity of glioma transcriptional cell states. Nat Genet (2021) doi:10.1038/s41588-021-00927-7.

40. Burr, M. L. et al. An Evolutionarily Conserved Function of Polycomb Silences the MHC Class I Antigen Presentation Pathway and Enables Immune Evasion in Cancer. Cancer Cell S1535610819303769 (2019) doi:10.1016/j.ccell.2019.08.008.

